# Hydrodynamics of the frontal strike in aquatic snakes: drag, added mass and the possible consequences for prey capture success

**DOI:** 10.1101/411850

**Authors:** Marion Segall, Anthony Herrel, Ramiro Godoy-Diana

## Abstract

Natural selection favors organisms that are the most successful in fitness-related behaviors such as foraging. Secondary adaptations pose the problem of re-adapting an already ‘optimized’ phenotype to new constraints. When animals forage underwater, they face strong physical constraints, particularly when capturing prey. Successful prey capture requires a predator to be fast and to generate a high acceleration. This involves two main constraints due to the surrounding fluid: drag and added mass. Both constraints are related to the shape of the animal. We experimentally explore the relationship between shape and performance in the context of an aquatic snake strike. As a model, we use two different 3D- printed snake heads representing typical shapes of aquatically-foraging and non-aquatically- foraging snakes, and frontal strike kinematics based on *in vivo* observations. By using direct force measurements, we compare the drag and added mass faced by the aquatic and non-aquatic snake models during a strike. Our results show that both drag and added mass are optimized in aquatic snakes. Using flow field measurements with particle image velocimetry, we examine the fluid dynamical mechanisms that could be behind the reduction of hydrodynamic constraints observed for the aquatic snake head shape, which makes it well suited to capture prey under water.

**Summary statement:** The present work explores the functional implications of head shape 15 in a group of aquatic predators using a fluid mechanics approach.

## Introduction

Aquatic animals have to overcome the strong viscous and inertial constraints associated with underwater movement ^1^. Physically, these constraints are related to the kinematics of movement and the morphology of an animal (i.e. the shape of the object that is facing the flow). For most aquatic vertebrates, viscous effects are confined to a thin boundary layer surrounding the body, which couples the motion of the animal with that of the surrounding fluid and gives rise to the skin friction that penalizes aquatic locomotion. In addition, fluid inertia causes the boundary layer to separate from the animal’s body, creating the recirculation zones associated to pressure drag ^2^. The specifics of the flow separation determine the relative importance of pressure to skin friction drag ^3,4^. In addition to drag, which depends on the velocity of the animal, the hydrodynamics are also dependent on acceleration of the added mass ^5,6^. This corresponds to the mass of fluid that is accelerated together with the animal and which exerts a reaction force. Both drag and added mass depend on the size and shape of the body ^5^, and it can thus be expected that the morphology of aquatic animals has evolved to reduce drag and added mass. However, organisms have a morphology that is also constrained by evolutionary history, functional trade-offs, and developmental programs thus restricting the range of possible morphological adaptations. Environmental and biological constraints act simultaneously on an organism and may all impact their evolution, sometimes leading to convergent phenotypes ^7–10^. Morphological convergence is common across the animal kingdom, yet its impact on function has only rarely been tested ^11–15^. We here use the case of convergence in head shape in aquatic snakes^16^ to provide an experimental test of the suggested functional advantages of observed similarities in the head shape of aquatic snakes.

Snakes are an ideal model to study convergence as they have invaded the aquatic medium multiple times independently throughout their evolutionary history. However, they do not show any of the usual adaptations to aquatic prey capture (e.g. they cannot perform suction feeding due to their reduced hyoid ^14^). Snakes have to deal with the hydrodynamic constraints when capturing a prey, and as these constraints are related with the shape ^1,13,17^, the head of aquatically foraging snakes should have evolved in a way to minimize the constraints. Convergence in head shape in aquatic snakes has been demonstrated previously ^14–16,18,19^. In a previous work ^16^, we compared the head shape of 62 species of snakes that capture prey under water (from sea snakes over homalopsids to North American watersnakes) versus 21 phylogenetically closely related species that do not forage under water. We used 3D geometric morphometrics on surface scans of these species and ran phylogenetic analyses demonstrating a morphological convergence in head shape of aquatically foraging snakes. Moreover, we characterized the shapes that are specific of both group of snakes (i.e. the aquatic and the non-aquatic foragers). We hypothesized that the convergent shape would provide a hydrodynamic advantage to aquatic foragers in comparison with their close relatives that do not capture aquatic prey. Several previous studies similarly have suggested convergence to give rise to a functional advantage ^13,14,16,20^, yet this has never been tested experimentally. Thus, we here propose an experiment to test this idea. In other words, we investigate whether the head shape associated with aquatically foraging snakes has a hydrodynamic advantage over the shape associated with the non- aquatic foragers. The hydrodynamic constraints involved during a strike are the pressure drag – skin friction being negligible in the regimes of interest here ^11^ – and the added mass. Both of these constraints are related to a certain extent to the shape of the object that is moving through a fluid ^5,6^. Thus, if our hypothesis is correct, the shape corresponding to the aquatic forager should show less drag and added mass than the non-aquatic model.

Another constraint related to the capture of prey under water is the mechanosensitivity of aquatic prey like fish. The lateral line system of fish is composed of mechanoreceptors that can detect very small pressure variations with an estimated threshold of 0.1 to 1 mPa at 1 mm ^21,22^. This system triggers a reflex escape response in the prey once a pressure threshold has been reached. Previous studies have suggested that a snake moving underwater generates a bow wave that might be able to trigger the reflex response of the prey ^11,14^. We tested this hypothesis and predicted that aquatic snakes should be stealthier than non-aquatic snakes during the strike such that the detection of the predator by the prey would be delayed.

We use direct force measurements on two 3D printed models of snake heads derived from our previous work based on the comparison of 83 species of snakes ^16^ (i.e. more than 400 snake specimens). As these models results from a 3D geometric morphometric analysis, the models are scaled to the same size, allowing us to specifically test for the impact of shape on hydrodynamic constraints. Our experimental setup mimics a ‘sit-and-wait’ frontal strike under water, meaning that the model remains motionless before the strike and is then suddenly accelerated to reach an almost constant speed for a short time. We compared models with the mouth open, as aquatic snakes keep their mouth opened during frontal strikes (Fabre et al., 2016; van Netten, 2006; Vincent et al., 2009, Herrel and Segall pers obs.). The force applied to the head during the strike was recorded to characterize the added mass and drag, which determine the hydrodynamic efficiency of a strike. In addition, another sensor was placed at the end of the strike track to assess the distance at which a prey is likely to detect the presence of the snake during capture. Particle Image Velocimetry (PIV) was used to visualize the flow field around the head during a strike. We also characterized the evolution of the vortex intensity during a strike for each shape, as it is closely related to the hydrodynamic forces generated by a moving object ^23–25^.

## Material & Methods

### 3D models

We compared two models that we termed “aquatic” and “non-aquatic” (Fig. 1). These shapes result from a 3D geometric morphometric study showing that the head shape of aquatic snake species has converged, possibly in response to the hydrodynamic constraints involved during prey capture under water ^16^. We compared the hydrodynamic forces that are exerted on each of the head shapes during a simulated capture event. The geometric morphometric analysis allows to extract shapes independent of variation of size such that the shapes are directly comparable to one another. In a next step we opened the mouth of the models as snake use to attack their prey with the mouth open. We used Blender^(tm)^ to rotate the jaw and the top of the head to reach an angle of 70^°^ based on previously published data on frontal strikes in snakes ^14,26,27^. The two models were then 3D printed using a Stratasys Fortus 250 MC 3D printer with ABS P430 as a material (Fig. 2a.).

**Figure 1:**
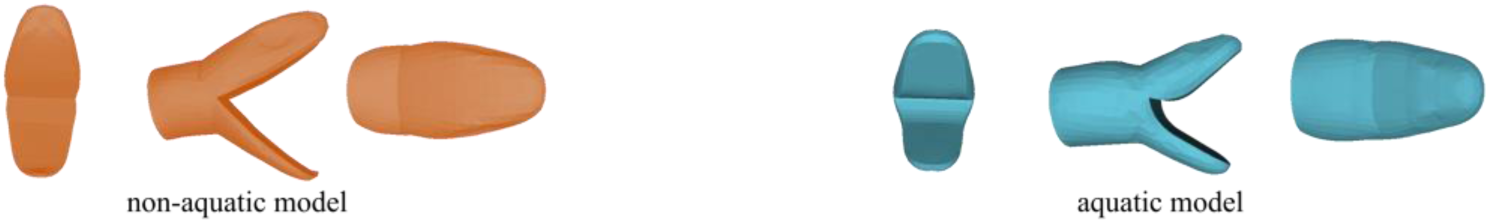
3D models of the head shape of non-aquatic (left) versus aquatic snakes (right) in front, side and top view.

**Figure 2:**
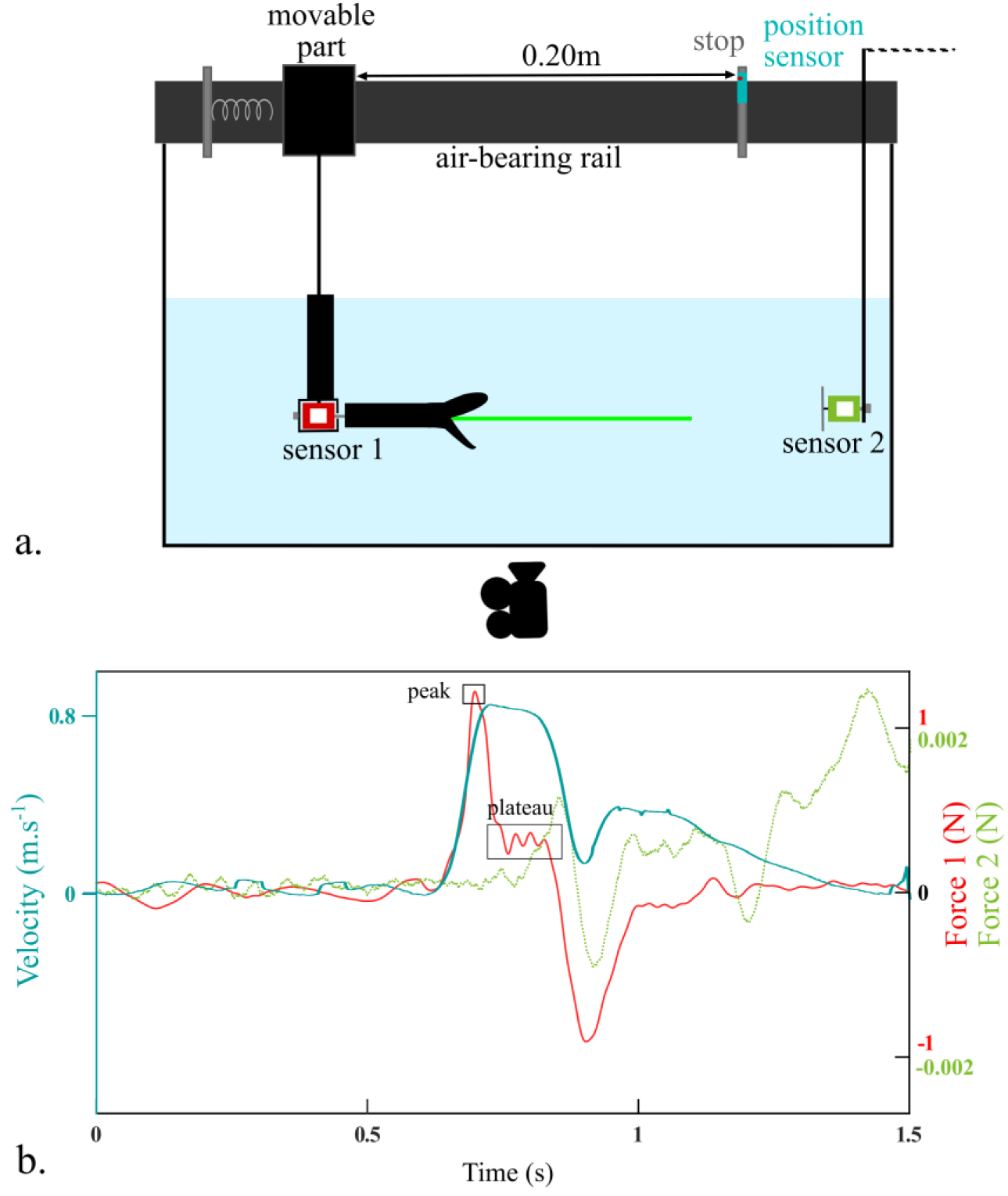
a. Experimental setup used to simulate a frontal attack of a snake towards a prey. ***b.*** Example of the output of the force sensor 1 (red line), force sensor 2 (green dashed line) and velocity (blue line) during one trial (i.e. one strike). The plateau and peak force used to calculate the hydrodynamic forces are indicated.

### Experimental setup

Snakes capture their prey using high acceleration forward motions that we mimicked using springs (Fig. 2a). We generated a range of speeds and accelerations by applying a different compression on the spring. We used a force sensor FUTEK LSB210+/-2 Lb to record the force exerted on the models which were positioned horizontally inside a water tank. This sensor was attached to the model using an aluminum rod and recorded the axial forces applied to the head during a strike. The other side of the sensor was attached to a bracket (sensor 1, Fig. 2a) that was itself hooked on the movable part of an air-bearing rail that allows the system to remain frictionless. This movable part was compressed against the spring and suddenly released. The length of the path was 20cm. Approximately 60 trials (i.e. spring compressions) were done for each model. To obtain the kinematics of each strike, we recorded the position of the movable part using a position sensor (optoNCDT1420, Micro-Epsilon) (Fig. 2a).

In addition, we wanted to assess what a prey would sense in terms of pressure, so we placed another, more sensitive, force sensor (FUTEK LSB210 100 g) at the end of the path to which we attached a round plastic piece of diameter 7 cm that allowed us to record the pressure changes (sensor 2, Fig. 2a). This sensor provided information about the distance at which a prey could potentially detect the presence of a snake during a strike. The force and position sensors were synchronized, and data were recorded at 1 kHz.

### Drag coefficient and added mass

The first part of the strike is the acceleration phase during which the velocity increases. This phase corresponds to the decompression of the spring. It is correlated with a dramatic increase in the force that is applied to the snake head model (red line, sensor 1, Fig. 2). Once the springs are completely decompressed, the system is no longer accelerating, and the velocity decreases slowly. In parallel, the force applied to the model decreases until it reaches a plateau-like phase (Fig. 2b). Then, the system hits the stop at the end of the track and moves backward generating a large drop in both velocity and force signaling the end of the trial.

During the plateau phase (Fig. 2b), the only force that is applied to the model and thus, the only force that is recorded by the sensor is the drag force. Thus, we used the average force recorded during this phase (F_d_) to calculate the drag coefficient (C_d_) of both of our models by using the standard definition ^2^:

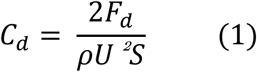

where *F_d_* is the drag force, *ρ* is the density of water, *U* the velocity of the object and *S* its projected frontal surface area, which was measured at 12.89 cm^2^ for the aquatic model and 14.72 cm^2^ for the non-aquatic model. The term 2*F_d_*/*ρS* was plotted against *U*^2^ and the linear regression coefficient corresponds to the drag coefficient of the models (Fig. 3). The Reynolds number range of our experiments is 1.10^4^-7.10^4^.

**Figure 3:**
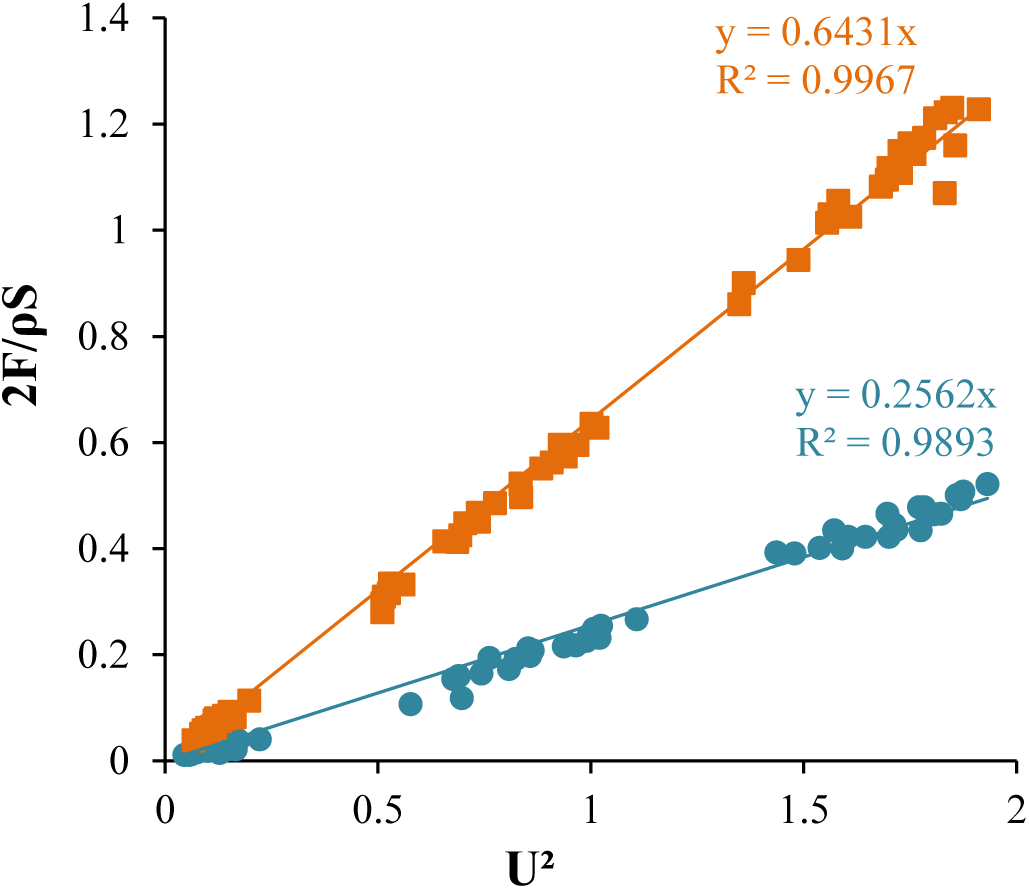
Drag term 2F_d_/ρS depending on the velocity term of the strike (U^2^) for the two head models tested. Linear regression lines are drawn. The slopes correspond to the drag coefficient of each shape and the R^2^ are the regression coefficients. Squares: non-aquatic model, circles: aquatic model.

During the acceleration phase, both drag and inertial forces are at play, meaning that the peak force (F_peak_, Fig. 2b) recorded by the force sensor is composed of these two forces. To calculate the added mass generated by both models, we used the following calculation steps for each trial, we first calculated the inertial force by subtracting the instantaneous drag force from the peak force measured by the sensor:

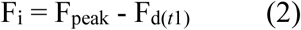

where F_i_ is the inertial force applied to the model and F_d(t)_ is the instantaneous drag force when the acceleration reaches its maximum:

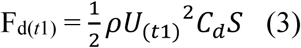

Here *ρ* is the density of water, U_(t1)_ the velocity at the instant the acceleration is maximal and *S* the projected frontal surface area of each model. Now, the added mass *M* can be computed as:

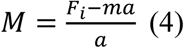

where *m* is the mass of the object, and *a* the acceleration.

Finally, the added mass coefficient (C_a_) ^2^:

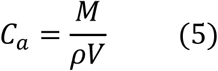

where *V* is the volume of the model: 7.33.10^-5^ m^3^ for the aquatic model and 5.78.10^-5^ m^3^ for the non-aquatic model.

The added mass coefficient was obtained by plotting the added mass term (*F_i_* - *ma*)/*ρV*, against the acceleration (a). The linear regression coefficient corresponds to the added mass coefficient of the models (Fig. 4).

**Figure 4:**
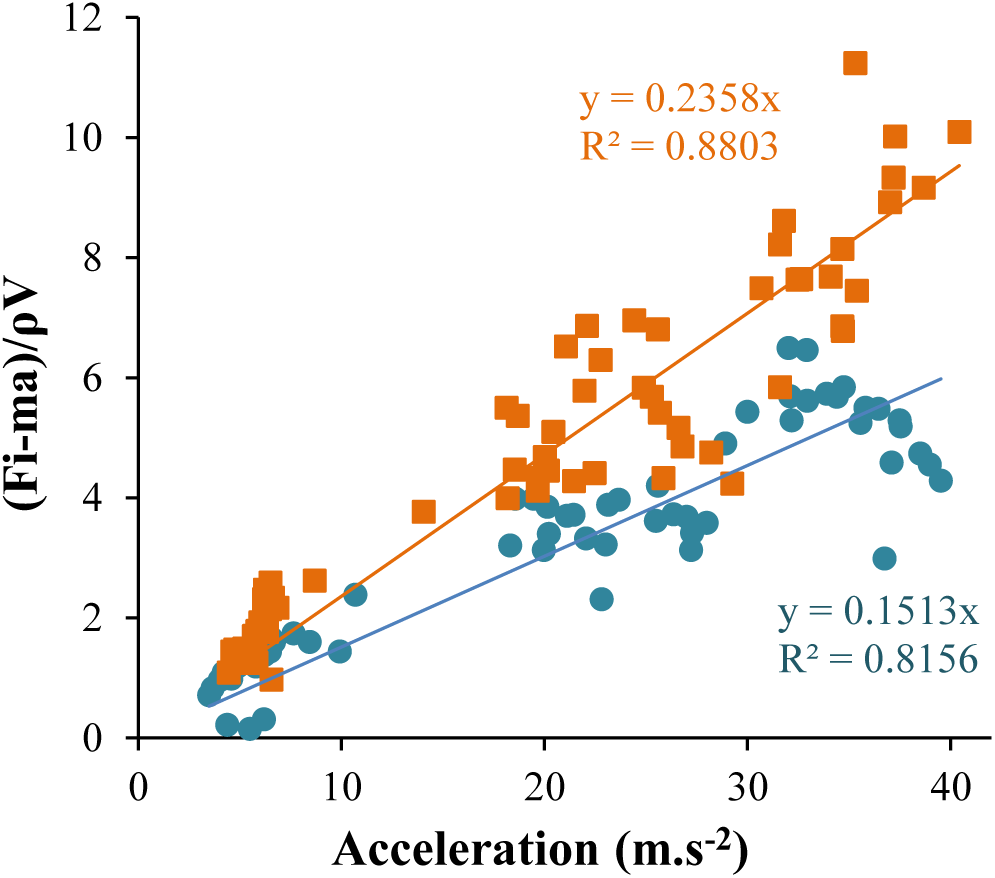
Normalized inertial force term (F_i_-ma)/ρV depending on the acceleration of the strike (a in m.s^-2^) for the two head models tested. Linear regression lines are drawn. The slopes correspond to the added mass coefficient of each shape and the R^2^ are the regression coefficients. Squares: non-aquatic model, circles: aquatic model.

### Detection distance

To compare the effect of the head shape on the detection by a possible prey we used the output of the second force sensor (sensor 2, Fig. 2a). This sensor can detect pressure variations of approximately 0.3 Pa which is around the hearing and the startle threshold of some fish (i.e. between 0.01Pa and 0.56Pa) ^28,29^. To estimate the position at which the prey could detect the predator, we defined the detection distance *d* as the position at which the force detected by sensor 2 deviates from the unperturbed value by more than one standard deviation of the sensor output before the strike (green dashed line, Fig 2b, Fig. 5).

**Figure 5:**
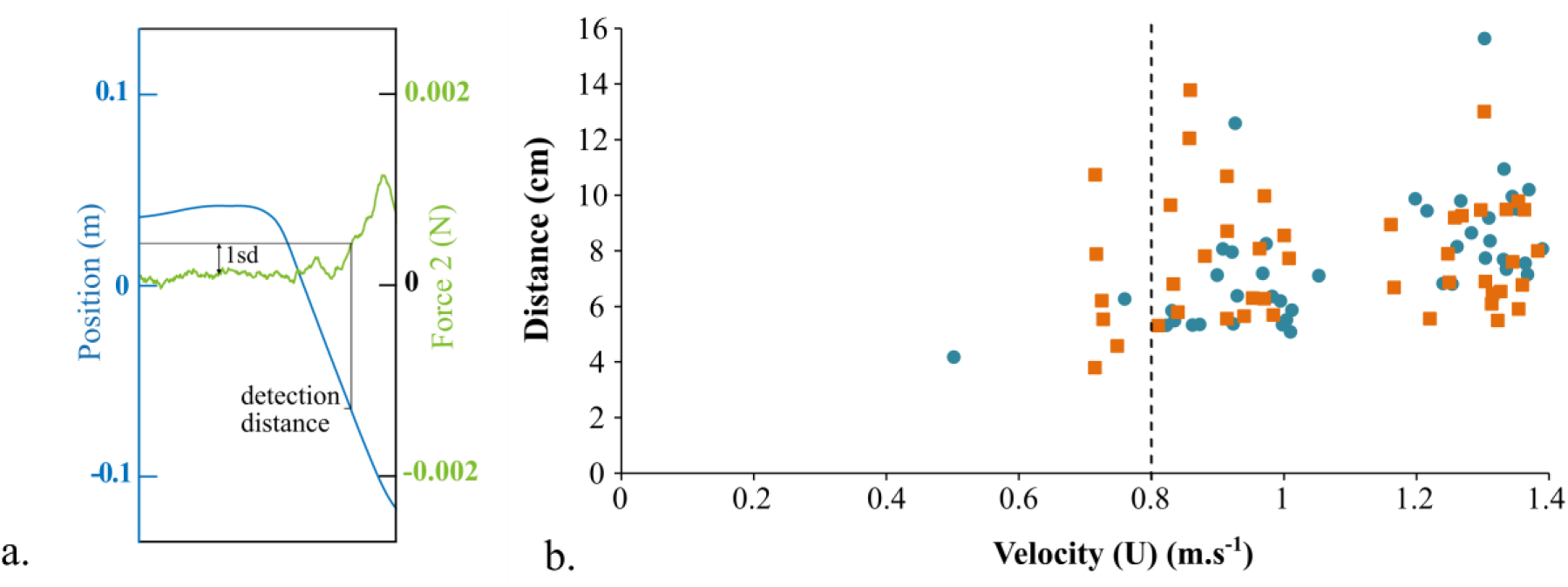
a. Zoom on the prey sensor output highlighting the method used to determine the detection distance, using the 1sd (standard deviation) threshold (not at scale here). ***b.*** Distance (cm) at which the prey could potentially detect the snake depending on the maximal velocity of the strike (m.s^-1^). For each graph: squares: non-aquatic model, circles: aquatic model.

### Particle Image Velocimetry

We used 2D Particle Image Velocimetry (PIV) with a high-speed camera, Dantec Dynamics SpeedSense M, to obtain a time-resolved recording of the strike from the bottom of the tank (Fig. 2a.). Water was seeded with polyamid particles of 20 μm in diameter and a Quantronix^®^ Darwin-Duo laser was used to produce the light sheet. Image acquisition was performed at 733Hz. We choose to record three different planes on each head to obtain a complete picture of the fluid flow around the head during the attack (see Supplementary Fig. S1). We applied the same compression to the springs (i.e. maximal compression) to get an equivalent comparison for the different shapes. Acquisition was performed using the Dantec DynamicStudio 2015a software. The PIV vector computation was performed using LaVision 7.2 with a 16 x 16 pixel^2^ interrogation window and 50% overlap. Additional post-processing and analysis was done in Matlab using the PIVMat toolbox ^30^. A more quantitative analysis was performed by computing the overall primary circulation *Γ* = ∫ *ω*^+^*dA* in each PIV plane (ω^+^ being the positive vorticity in Fig. 6b.). The evolution of the dimensionless circulation *Γ/UL* as a function of time, where L is the characteristic length scale of the acceleration regime of the strike maneuver (which is constant for all experiments) and *U* is the velocity of the strike is plotted in Fig. 6b.

**Figure 6:**
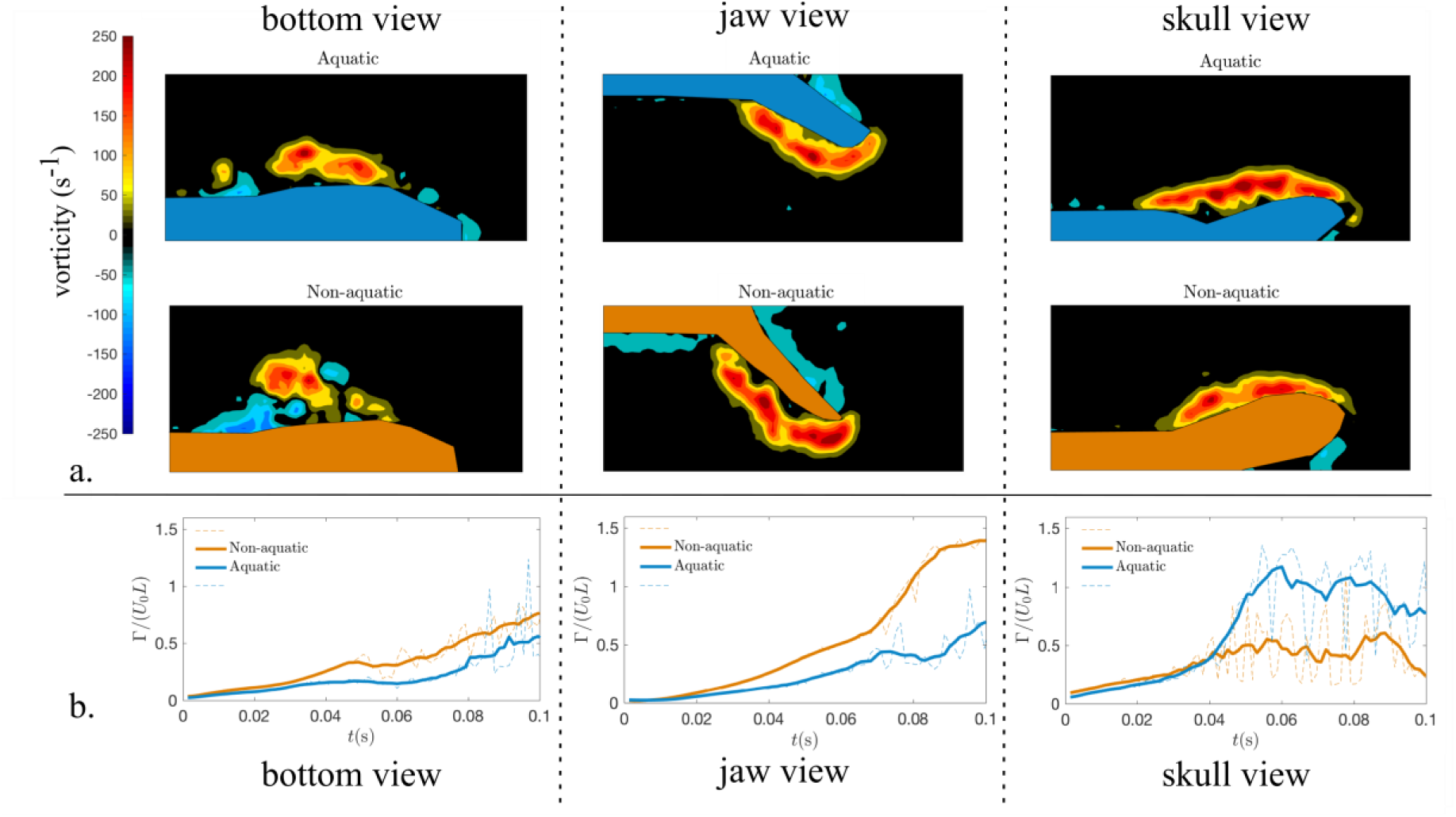
a. Snapshots of the vorticity field ω_z_ around the snake head models at the end of the acceleration phase for the aquatic (first line) and non-aquatic (second line) models, in the three measurement planes: bottom, jaw and skull views are shown on the first to third columns, respectively. The color bar for the vorticity field is given in s-1. ***b.*** Evolution of the dimensionless integrated positive circulation during the acceleration phase depending on the time for both models in each of the three views considered.

### Statistical analyses

To test for differences between the drag coefficients of the two shapes, we ran a Pearson correlation on the force component of the drag coefficient (*2F_d_/ρS*) with the square velocity (*U^2^*). An ANCOVA with mass as a co-variate was performed to test for statistical differences in the drag coefficient between the two models. To compare the detection distance, we ran an ANCOVA with the distance as the response variable, the model as a factor, and the velocity as covariate. All the variables were Log10-transformed and the statistical analyses were performed using R ^31^. The significance level was set at 5%.

## Results

### Drag and added mass

The drag coefficient of the non-aquatic shape is higher than the coefficient of the aquatic model, respectively 0.64 and 0.26 (Pearson’s correlation: nonaq: df = 67, *P* < 0.001, R^2^ = 0.996; aq: df=64, *P* < 0.001, R^2^ = 0.995; ANCOVA: *F*_2,132_ = 671.1, *P* < 0.001) (Fig. 3).

The mean added mass obtained is 12.67 g for the aquatic model versus 14.95 g for the non-aquatic model. The added mass coefficients obtained from the linear regression on Fig. 4 are 0.151 for the aquatic model and 0.235 for the non-aquatic model.

### Detection distance

The force signal was too noisy to get any accurate measures of the detection distance at low velocities (i.e. U > 0.5 m.s^-1^). There is moreover no statistical difference between the distances at which the prey could detect the presence of the snake depending on their head shape. However, this distance depends on the maximal velocity of the strike, the faster the strike, the earlier the detection of the predator (ANCOVA: *F*_2,84_ = 5.05; *P* = 0.008; model: *P* = 0.65; Umax: *P* = 0.008) (Fig. 5).

### Flow characterization

The frontal strike maneuver involves strong flow separations due to the high shear produced by the impulsive acceleration. The flow features can be characterized by examining the vortex structures formed at the corner of the mouth and on both tips of the jaw and of the skull. We created videos of the vortex formation during a strike, obtained from PIV in three planes around the snake heads (see Materials and Methods section), to compare both models (see Supplementary videos S2-4). The PIV measurements show the formation of vortices during the strike maneuver. In Fig. 6a, we compare the vorticity field at the end of the acceleration phase (at t ≈ 0.8 s) in the three measurement planes; bottom view, jaw view, and skull view (Supplementary Fig. S1) for the aquatic and non-aquatic heads. Looking at the bottom view, the advantage of the aquatic model seems to be related to a smaller primary vortex. The picture is not as straightforward considering the jaw and skull view, where opposite observations on the primary vorticity production can be observed qualitatively: on the jaw view the primary vorticity patch appears more detached from the jaw in the non-aquatic case, whereas in the skull view the same is true for the aquatic case. Fig. 6b shows the quantitative analysis of the primary circulation. First, we can see that in the bottom view the aquatic model induces a slightly (∼10%) lower overall circulation over the whole acceleration phase. Second, for the jaw view it can be remarked that a much lower overall circulation is produced by the vorticity detached from the tip of the jaw in the aquatic case (around 40% of the non-aquatic value at the end of the acceleration phase). The picture in the skull view is the opposite with the aquatic shape generating more overall circulation but the difference between the two models is less important than for the jaw view. We note also for the skull view that the computed value for the circulation is more variable.

## Discussion

Drag is well known for its importance during steady locomotion. However, it is also involved in transient behaviors such as the capture maneuver studied here. Certainly, the aquatic shape appears better adapted to capture aquatic prey using a frontal strike than the non-aquatic shape in terms of drag. The aquatic model has a drag coefficient that is almost 3 times smaller than the non-aquatic model. As mentioned above, drag in this fast-impulsive maneuver is mainly pressure drag, which is intimately linked to the flow separation in the near wake of the snake head as it moves. The PIV measurements illustrate the vortices that are formed very early during the strike (see Supplementary videos S2-4). Looking at the bottom view in Fig. 6, the drag advantage of the aquatic model could be related to a smaller primary vortex, the non-aquatic case showing a more fluctuating and disordered flow field. Moreover, the vorticity produced at the tip of the jaw shows a clear quantitative difference and is consistently higher for the non-aquatic model. However, the skull view shows the opposite pattern of vorticity; the non-aquatic shape produces fewer vortices with an integrated primary circulation that is less important than for the aquatic model. It should be noted that the 2D nature of the PIV measurements presented here does not allow us to provide a quantitative link between drag and the vorticity profile of the flow around the head. Nonetheless, from the present results we can conjecture that a reduction of the recirculation bubble behind the jaw may be one of the main physical mechanisms explaining the physical advantage of the head shape observed in aquatically foraging snakes.

Transient maneuvers under water, such as the underwater prey capture in snakes, implicate an acceleration phase that not only involves drag but also inertia. Inertial forces under water are associated with the mass of the object but also with a mass of the fluid that is accelerated. Thus, the relationship between inertia and shape is not straightforward. However, some studies suggested that an optimal body shape for transient propulsion, such as a snake strike, would be an elongated, streamlined, and flexible body and non-muscle mass reduction, which corresponds to a snake-like configuration ^1,17^. To our knowledge, no study to date has focused on the shape of the head and its role. In this study, we highlight that the hydrodynamic forces associated with a transient maneuver are important in comparison with drag (e.g. the peak of force in comparison with the plateau on Fig. 2). Moreover, we demonstrated that the aquatic shape allows to reduce the added mass and is associated with a smaller added mass coefficient. This suggests that drag is not the only driver of the evolution of head shape in aquatic snakes. Moreover, added mass and drag optimization do not require divergent morphological features in the case of aquatic snake strikes, unlike what suggested for the body shape of fish ^17^.

Regarding the prey detection distance, our results show that this distance does not depend on the snake head shape, but rather that it increases with strike velocity. However, we cannot conclude on the biological relevance of the absolute prey detection distance measured in our experiment as our setup was built with as primary purpose to measure drag and added mass. Snakes usually strike when the prey is close to their head (e.g. 0.5-0.8 cm for *Erpeton tentaculatum* ^32^; 4.87 cm for *T. couchii*; 2.81 cm for *T rufipunctatus* ^33^; less than 3 cm for *Hydrophis schistosus* ^34^). The detection distance measured here is around 6 to 10 cm, so we could consider that the prey can possibly detect the snake almost instantaneously upon the strike initiation, the reaction time of a fish being around 7 ms ^32^. Capture success is thus more likely determined by the hydrodynamic profile of the snake head than being dependent on the reaction of the prey.

In conclusion, we investigated the role of head shape on the hydrodynamic forces generated by a predator using an experimental approach focusing on a transient maneuver. We were able here to quantify the role and impact of head shape in the hydrodynamics of prey capture in aquatic snakes. We highlighted a clear hydrodynamic advantage of the aquatic head shape when capturing a prey being associated not only with a smaller drag coefficient but also a smaller added mass coefficient. These results validate the hypothesis that the morphological convergence of the head shape in aquatic snakes is an adaptation to an aquatic lifestyle as it provides a clear hydrodynamic advantage. In this work, we focused on the shape of the head of aquatically foraging snakes, as several studies have highlighted convergence therein, and as shape plays a crucial role in the hydrodynamic constraints as well. Size could be an important feature regarding the hydrodynamic constraints. However, we did not detect any allometry in our morphological study, meaning that the aquatically foraging snakes are not significantly different in size than their closely related non-aquatic species. Thus, the present work focuses on the functional meaning of shape irrespective of size. The other factors that could play a role in the hydrodynamics of the prey capture of aquatic snakes could be the gape angle and macro and microscopic skin features which remains to be investigated.

## Acknowledgments

We thank Olivier Brouard, Amaury Fourgeaud and Tahar Amorri from the PMMH lab for their precious help in the experimental design as well as Xavier Benoit-Gonin for his help with the 3D printer. Thierry Darnige and especially Justine Laurent are acknowledged for their help with the sensors and computer coding. MS thanks the Région Ile de France for funding this research project and the doctoral school Frontières du Vivant (FdV) – Programme Bettencourt.

## Author contributions

All authors helped revise and approved the manuscript and conceived the study. MS carried out the data collection, the statistical analyses, and wrote the manuscript. RGD helped to build the experimental setup and to interpret the data. RGD carried out the particle image velocimetry analysis. AH participated in the scientific interpretation of the data in a biological context.

## Data availability

See Supplementary Table S5

